# Room-temperature serial synchrotron crystallography of apo PTP1B

**DOI:** 10.1101/2022.07.28.501725

**Authors:** Shivani Sharma, Ali Ebrahim, Daniel A. Keedy

## Abstract

Room-temperature X-ray crystallography provides unique insights into protein conformational heterogeneity, but a common hurdle is obtaining sufficiently large protein crystals. Serial synchrotron crystallography (SSX) helps address this hurdle by allowing the use of many medium- to small-sized crystals. We have used a recently introduced serial sample support chip system to obtain the first SSX structure of a human phosphatase, specifically Protein Tyrosine Phosphatase 1B (PTP1B) in the unliganded (apo) state. In previous apo room-temperature structures, the active site and allosteric sites adopted alternate conformations, including open and closed conformations for the active-site WPD loop and for a distal allosteric site. By contrast, in our SSX structure, the active site is best fit with a single conformation, but the distal allosteric site is best fit with alternate conformations. This observation argues for additional nuance in interpreting the nature of allosteric coupling in this protein. Overall, our results illustrate the promise of serial methods for room-temperature crystallography, as well as future avant-garde crystallography experiments, for PTP1B and other proteins.

## Introduction

X-ray crystallography offers detailed insights into protein structure. Although most X-ray crystallography is performed with the sample at cryogenic temperature (cryo), data collection at room temperature (RT) offers unique insights into protein conformational heterogeneity and function (Fraser et al. 2009, 2011; Keedy et al. 2014, 2015; Fischer, Shoichet, and Fraser 2015; Milano et al. 2022). Previously, a barrier to RT crystallography was growing sufficiently large crystals to counteract the increased sensitivity to radiation damage at RT relative to cryo (Garman and Nave 2009; Warkentin et al. 2013). However, new methods are emerging to enable RT crystallography (Fischer 2021). In particular, serial crystallography methods which enable the use of hundreds to thousands of microcrystals instead of larger macrocrystals have been developed for X-ray free electron lasers (XFELs) (Hirata et al. 2014; Moreno-Chicano et al. 2019), and have also been applied at synchrotrons in an approach known as serial synchrotron crystallography (SSX) (Owen et al. 2017; Diederichs and Wang 2017).

Recently, a new serial sample support system for SSX was introduced (Illava et al. 2021). With this approach, a vacuum source in a humid chamber is used to load microcrystals onto a chip, which has a base that is compatible with standard goniometers at synchrotron beamlines. The system can be used at cryo, but is particularly valuable for RT. It has been previously demonstrated for three proteins: fluoroacetate dehalogenase, hen egg-white lysozyme, and human glutaminase C (Illava et al. 2021; Milano et al. 2022).

Here, we exploit this serial system for a distinct, biomedically relevant protein target: human Protein Tyrosine Phosphatase 1B, or PTP1B (also known as PTPN1). PTP1B is the archetypal PTP, and is a validated therapeutic target (S. Zhang and Zhang 2007) for diabetes (Z.-Y. Zhang and Lee 2003; Montalibet and Kennedy 2005), cancer (Tonks and Muthuswamy 2007), Alzheimer’s disease (Vieira et al. 2017), and Rett syndrome (Krishnan et al. 2015). Because active-site inhibitors of PTP1B suffer from bioavailability and selectivity limitations (Stanford and Bottini 2017), notwithstanding some progress in bypassing such limitations (Z. Y. Zhang 2001), there has been increasing focus on the potential of allosteric inhibition. Several allosteric inhibitors targeting different sites in PTP1B have been reported in the literature (Wiesmann et al. 2004; Hansen et al. 2005; Krishnan et al. 2014, 2018; Keedy et al. 2018; Friedman et al. 2022). However, to date, none have been clinically approved, illustrating the persistent need to elucidate conformational ensembles and allosteric mechanisms in this protein.

As noted above, RT crystallography can reveal protein alternate conformations, which can interact with one another to imbue proteins with allosteric properties (van den Bedem et al. 2013). For PTP1B, 295 cryo structures are available in the Protein Data Bank (PDB) (Berman et al. 2000), but only 7 RT structures are available, only 4 of which are of the apo enzyme. The first 2 such structures (PDB ID 6b8x and 6b8t) are from multitemperature crystallography of PTP1B that indicated the existence of an extensive allosteric network spanning several promising sites in the protein (Keedy et al. 2018). Although those 2 structures are nominally apo, the crystals included high concentrations of glycerol, resulting in glycerol molecules that bound in the active site and ostensibly biased the protein’s conformational ensemble (Keedy et al. 2018). Of the remaining 2 RT structures of PTP1B, 1 is in the same crystal lattice as 6b8x but has not yet been described in the literature; the other (PDB ID 2cm2) is in a space group with more extensive crystal contacts to key sites, has a partially ordered 6xHis expression tag in an allosteric site that may bias the conformational ensemble there and elsewhere, and has no structure factors available. No serial crystallography structures of PTP1B-nor indeed, to our knowledge, of any human phosphatase -- are yet available.

Here, we provide the first serial synchrotron crystallography (SSX) structure of PTP1B, demonstrating this method’s feasibility for this protein. We use two different processing pipelines and obtain very similar results with both, demonstrating the robustness of processing data from the SSX chips used here. Our dataset allows us to draw comparisons to other existing structures with regards to the conformational ensemble of truly apo PTP1B. As may be expected, our model is broadly similar to the dozens of ligand-bound cryo structures, and the smaller set of RT structures in different conditions. However, despite its moderate resolution relative to past single-crystal structures, our dataset also indicates a degree of allosteric decoupling that adds nuance to the previously reported paradigm of allostery in this protein (Keedy et al. 2018). Finally, our experiments pave the way to future serial crystallography experiments for PTP1B and related proteins.

## Results

To determine a room-temperature serial synchrotron crystallography structure of PTP1B, we used a recently introduced fixed-target serial sample support system, including multifaceted chips that are loaded with microcrystals in a custom humidified environment (Illava et al. 2021) (**Supp. Fig. 1**). Using this system at the MacCHESS ID7B2 beamline, we loaded 6 chips with PTP1B crystals in the previously characterized P 31 2 1 space group (Pedersen et al. 2004; Keedy et al. 2018). Across these chips, we collected a total of 1297 partial datasets (wedges) constituting 1.2-3.0° each. The diffraction weighted dose for each crystal in this experiment was estimated using RADDOSE-3D (Bury et al. 2018) to be < 35.1 kGy per crystal (see Methods), suggesting an absence of substantial radiation damage.

To ensure our data were processed robustly, we used two pipelines in parallel, XDS (Kabsch 2010) and DIALS (Winter et al. 2022), and compared the results. The two pipelines involved different software, but had similar overall logic: splitting into wedges; spot-finding, indexing, and integration for each wedge; and scaling and merging across all wedges (**Fig. 1**). Both pipelines yielded similar numbers of successfully processed wedges across the 6 chips (**Supp. Table 1**) and similar overall statistics after scaling and merging (**Table 1**).

**Figure 1:**
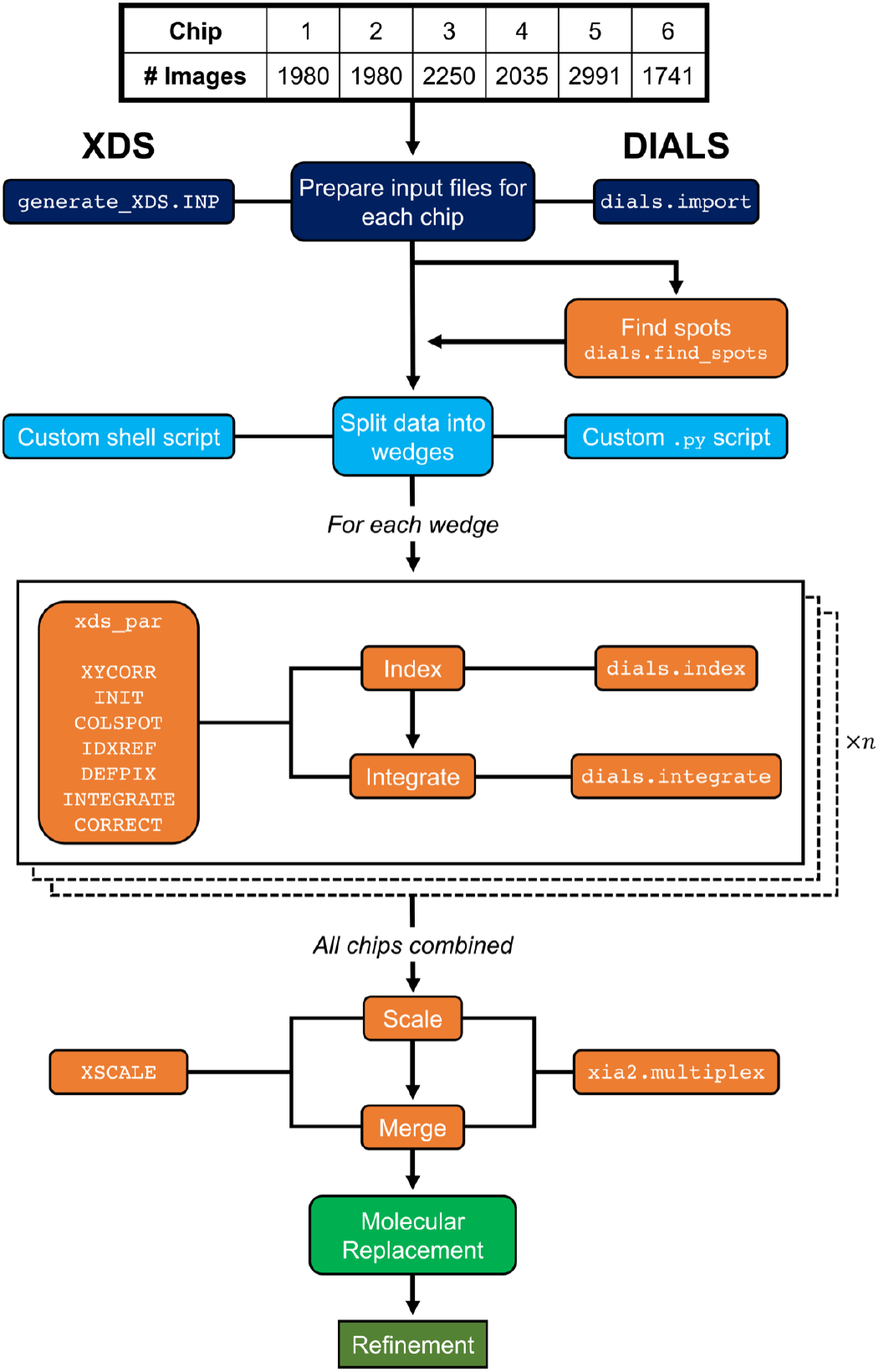
Flowchart of data reduction workflow. Two pipelines, XDS and DIALS, were used in parallel to process our SSX data. Both involved similar workflows (central flow), albeit with slight variations (left vs. right). *‘ní* refers to the number of wedges.

**Table 1:** Crystallographic statistics for XDS and DIALS data reduction. Overall statistics given first (statistics for highest-resolution bin in parentheses).

| Method | XDS | DIALS |
| --- | --- | --- |
| Resolution (Å) | 44.67-2.40 | 105.5-2.40 |
| Completeness (%) | 99.7 (99.7) | 99.82 (100.0) |
| Multiplicity | 11.45 (11.06) | 12.10 (12.68) |
| I/sigma(I) | 4.62 (1.08) | 4.5 (0.8) |
| R<sub>merge</sub>(I) | 0.661 (4.23) | 0.938 (5.921) |
| R<sub>meas</sub>(I) | 0.693 (4.754) | 0.980 (6.166) |
| R<sub>pim</sub>(I) | 0.201 (1.307) | 0.272 (1.666) |
| CC<sub>1/2</sub> | 0.881 (0.135) | 0.929 (0.218) |
| Wilson B-factor (Å<sup>2</sup>) | 47.57 | 40.81 |
| Total observations | 223455 (15663) | 234336 (11935) |
| Unique observations | 19515 (1416) | 19359 (941) |
| Space group | P 31 2 1 | P 31 2 1 |
| Unit cell dimensions<br>(Å, Å, Å, °, °, °) | 89.35, 89.35, 105.76,<br>90, 90, 120 | 89.06, 89.06, 105.48,<br>90, 90, 120 |

Therefore, we arbitrarily proceeded with XDS. The overall “hit rate” was ~10%, with 129 successfully processed wedges for XDS (similar for DIALS) out of 1297 total wedges collected. This efficiency could likely be improved with experimental optimization, which was not performed here. Analysis of correlation coefficients between all the unmerged wedges with XSCALE_ISOCLUSTER (Diederichs 2017) indicated the existence of only one cluster, thus obviating the need for merging separate isomorphous clusters of subsets of the data.

We used molecular replacement and iterative model building and refinement to obtain a final RT SSX structural model of apo PTP1B, which has suitable fit-to-data and model validation statistics (**Table 2**). Using this model, we next inspected the protein’s conformational ensemble at several key sites (**Fig. 2**).

**Table 2:** Crystallographic statistics for XDS data reduction and structural modeling. Overall statistics given first (statistics for highest-resolution bin in parentheses).

|  |  |
| --- | --- |
| PDB ID | 8du7 |
| Method | XDS |
| Resolution (Å) | 2.40 |
| Completeness (%) | 99.7 (99.7) |
| Multiplicity | 11.45 (11.06) |
| I/sigma(I) | 4.62 (1.08) |
| R<sub>merge</sub>(I) | 0.661 (4.23) |
| R<sub>meas</sub>(I) | 0.693 (4.754) |
| R<sub>pim</sub>(I) | 0.201 (1.307) |
| CC<sub>1/2</sub> | 0.881 (0.135) |
| Wilson B-factor (Å<sup>2</sup>) | 47.57 |
| Total observations | 223455 (15663) |
| Unique observations | 19515 (1416) |
| Space group | P 31 2 1 |
| Unit cell dimensions<br>(Å, Å, Å, °, °, °) | 89.35, 89.35, 105.76,<br>90, 90, 120 |
| Solvent content (%) | 62.64 |
| R<sub>work</sub> | 0.195 |
| R<sub>free</sub> | 0.237 |
| RMS bonds (Å) | 0.013 |
| RMS angles (°) | 1.33 |
| Ramachandran outliers (%) | 0.35 |
| Ramachandran favored (%) | 92.55 |
| Clashscore | 6.06 |
| MolProbity score | 2.12 |

**Figure 2:**
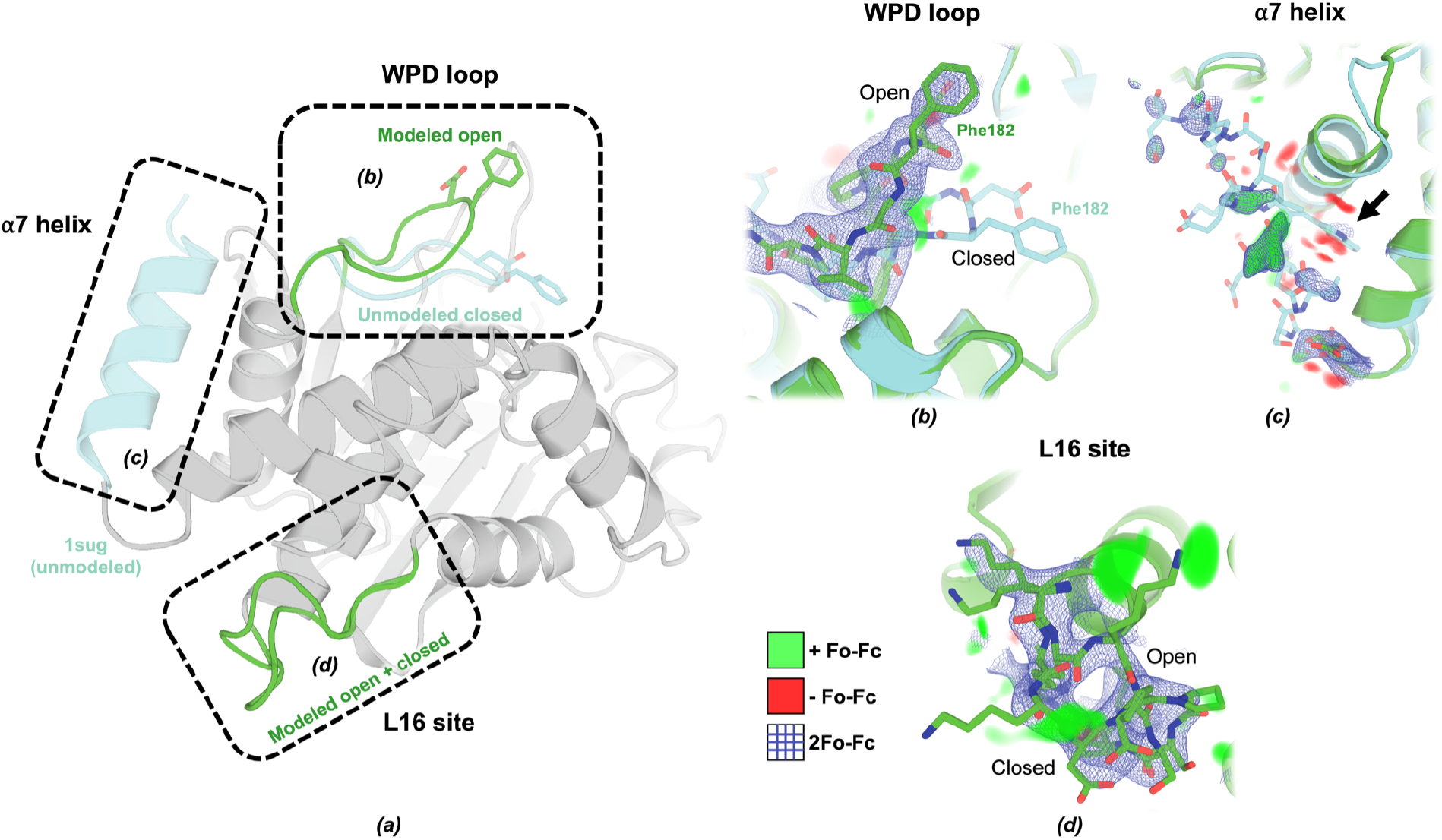
Conformational heterogeneity across the PTP1B allosteric network in an SSX RT structure. (*a*) Overview of key sites in PTP1B featured in (*b-d*). (*b*) The active-site WPD loop is best modeled as adopting only the open conformation, based on 2Fo-Fc electron density contoured at 1 σ (blue) and +/− Fo-Fc difference electron density contoured at +/− 3.0 σ (green/red). The closed conformation of the WPD loop as modeled in PDB ID 1sug (transparent cyan) is shown for comparison. See also **Supp. Fig. 2 (dual-conf)**. (*c*) The α7 helix is best modeled as disordered, based on 2Fo-Fc density at 1 σ and Fo-Fc density at +/− 3.0 σ. The ordered conformation of α7 from 1sug is shown for comparison. Note the absence of strong density for the Trp291 “anchor” that can occupy the allosteric BB site (arrow). See also **Supp. Fig. 3 (a7 refined)**. (*d*) In contrast to the WPD loop and α7, Loop 16 definitively adopts alternate conformations with similar occupancies, based on 2Fo-Fc density at 1 σ and Fo-Fc density at +/− 3.0 σ. See also **Supp. Fig. 4 (omits)**.

First, the active-site WPD loop adopts the open conformation (**Fig. 2b**), which is known to dominate in solution (Whittier, Hengge, and Loria 2013). There is some residual +Fo-Fc electron density below this open conformation near the location of the closed conformation seen in previous structures (Pedersen et al. 2004). However, refinement with a dual-conformation model as used previously (Keedy et al. 2018) resulted in an absence of 2Fo-Fc density above 0.7 σ for the closed conformation (**Supp. Fig. 2**), arguing against this interpretation of the data. Some residual density is detectable below the WPD loop and may be attributable to the complex water network of the active site (Pedersen et al. 2004), although the moderate resolution of our dataset limits our ability to interpret the details of this network.

A second key site is the α7 helix, which has been established by a variety of methods as a key allosteric hub in PTP1B (Olmez and Alakent 2011; Krishnan et al. 2014; Choy et al. 2017; Keedy et al. 2018) as well as its closest homolog, TCPTP (Singh et al. 2021). In our SSX structure of PTP1B, the α7 helix cannot be confidently modeled in the ordered conformation, and thus is best left disordered as in many previous open-state structures (**Fig. 2c**). An attempt at modeling the ordered conformation results in detectable but weak electron density support (**Supp. Fig. 3**) and inflated B-factors of >100 Å^2^ (as compared to ~40-60 Å^2^ for a more fully ordered α7 in PDB ID 1sug).

A third key site in the protein, Loop 16, is the eponymous loop of the reported allosteric L16 site (Keedy et al. 2018), which also involves the adjacent protein N-terminus and the C-terminus of the α6 helix as it transitions to α7. The L16 site constitutes a cryptic site (Cimermancic et al. 2016) that accommodates ligands only in its open conformation, as discovered in a high-throughput crystallographic small-molecule fragment screen (Keedy et al. 2018), adding to its potential value as a targetable allosteric site in PTP1B. In our SSX structure, unlike the WPD loop and α7 helix, PTP1B clearly adopts alternate conformations for Loop 16, for the open and closed states (**Fig. 2d**). These conformations have similar occupancy in the refined model (46% open, 54% closed), suggesting they are approximately isoenergetic in our experimental conditions. Omit maps of either conformation result in convincing difference density features, confirming the simultaneous presence of both states in the crystal (**Supp. Fig. 4**).

These observations from our SSX structure are in contrast to the previous report of an allosteric network in PTP1B with coupled opening of the WPD loop, opening of the L16 site, and disordering of the α7 helix (Keedy et al. 2018). That concept is embodied in PDB ID 6b8x (and 6b8t), which have open/closed alternate conformations for the WPD loop, open/closed alternate conformations for the L16 site, and a quasi-ordered partial-occupancy α7 (**Fig. 3**). Another recently deposited RT structure of apo PTP1B, PDB ID 7rin (no publication available), agrees with 6b8x in these respects. 6b8x has the same space group and unit cell dimensions as our SSX structure, so its more discernible conformational heterogeneity may instead be related to its higher resolution (1.74 Å as opposed to 2.40 Å here). However, PDB ID 2cm2 is even higher-resolution (1.50 Å), yet features a single-conformation open WPD loop, open L16 site, and disordered α7 (Ala et al. 2006) (**Fig. 3**). Thus high resolution is not sufficient to resolve these alternate conformations. Instead, the different space group, unit cell, and crystal contacts of 2cm2 may explain its collapsed conformational ensemble at these sites. Unfortunately, structure factors are not available for 2cm2, so we cannot interrogate its electron density map for signs of “hidden” unmodeled alternate conformations (Lang et al. 2010), as detected previously for PDB ID 1sug (Keedy et al. 2018). Overall, comparison of our SSX model with previous structures indicates partial decoupling of the allosteric network in PTP1B, suggesting that coupling within this network may not be as tight as originally envisioned.

**Figure 3:**
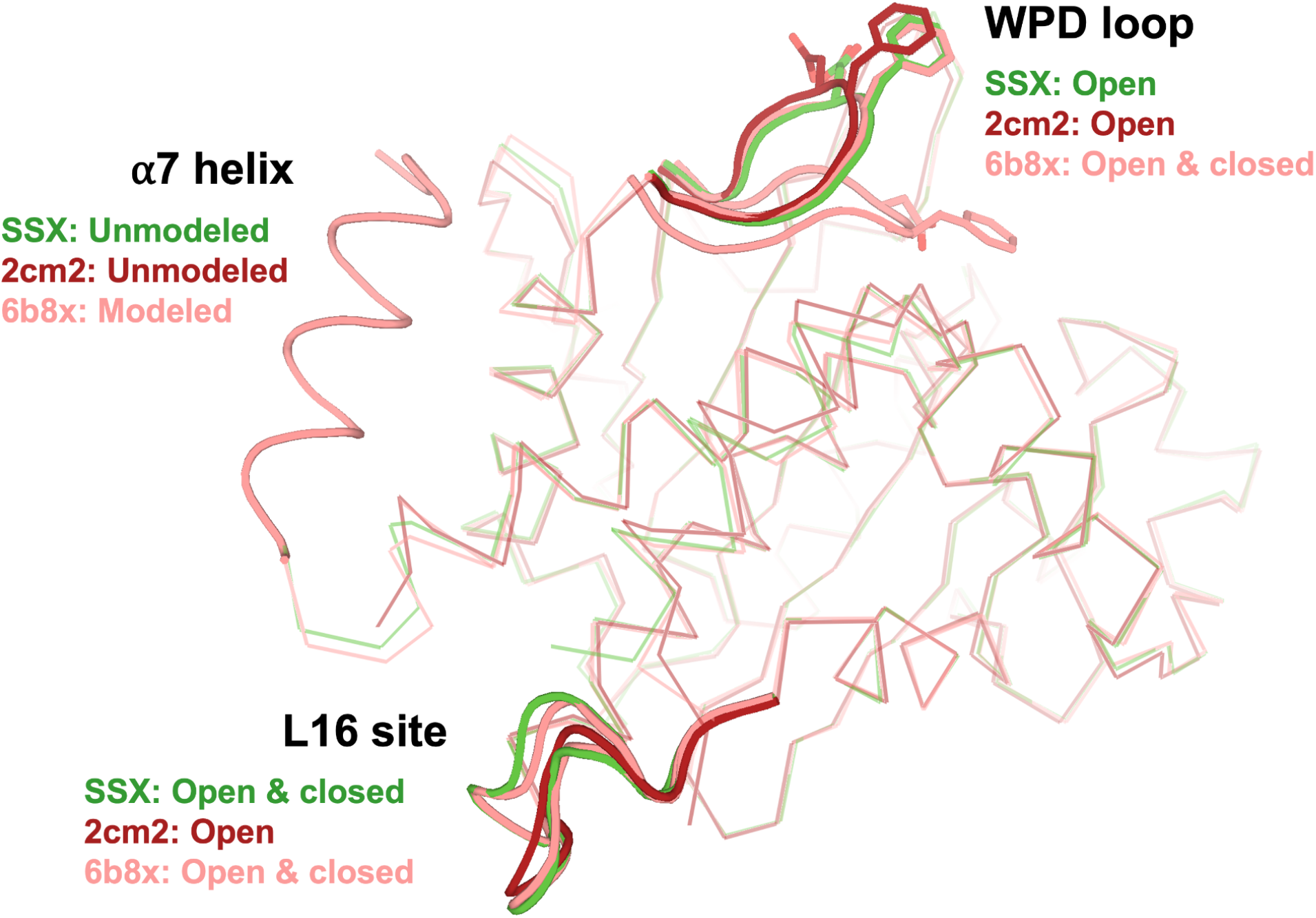
SSX RT structure vs. previous RT structures of PTP1B have different patterns of conformational heterogeneity across allosteric network. Overlaid are our SSX structure (green), PDB ID 2cm2 (red), and PDB ID 6b8x (pink).

## Discussion

Here we present the first SSX structure of a human phosphatase, in this case PTP1B. The serial sample support system used here (Illava et al. 2021) is flexible, allowing data collection at RT or cryo using standard X-ray beamline goniometers. It thus represents a positive addition to the expanding toolkit for RT crystallography experiments (Fischer 2021).

Unsurprisingly, our SSX structure of PTP1B is broadly similar to previous single-crystal structures in this and other space groups. Nevertheless, demonstrating the feasibility of serial experiments for this biomedically central protein is a valuable step. Moreover, our results add nuance to the previously reported picture of the allosteric network in PTP1B, in which opening of the WPD loop, disordering of the α7 helix, and opening of Loop 16 are highly correlated (Keedy et al. 2018). In the SSX structure, it appears that the expected coupling between the WPD loop and α7 is present: the former is open (**Fig. 2b**, **Supp. Fig. 2**) and the latter is disordered (**Fig. 2c**, **Supp. Fig. 3**). However, Loop 16 adopts both open and closed conformations (**Fig. 2d**), suggesting that the coupling between the L16 site and the other two areas may not be as tight as previously hypothesized. This reframing is reminiscent of how the active-site side-chain network of the proline isomerase CypA was previously viewed as thoroughly coupled (Fraser et al. 2009), but reinterpreted as involving hierarchical coupling on the basis of a multitemperature crystallography series (Keedy et al. 2015). Such observations highlight the need for multidataset approaches to crystallography and accompanying multiconformer modeling to better infer allosteric mechanisms in proteins (Keedy 2019).

Although we observe different coupling between sites in our SSX structure vs. previous RT apo structures, some caveats should be considered that complicate this interpretation. First, the resolution of our structure is only moderate (2.40 Å) when compared to previous single-crystal structures (<2 Å), although this could likely be improved by optimizing our crystals for serial diffraction, which we did not undertake. Second, although the unit cell and crystal contacts in our structure are the same as PDB ID 6b8x, they are different from 2cm2. However, moving forward, such crystal lattice differences can present opportunities to learn from differential localized perturbations to surface regions of proteins -- conceptually similar to how crystallographic alternate conformations were often viewed as a nuisance during model building and refinement, but can now be exploited to learn about biological function (Fraser et al. 2009).

In closing, we look forward to the ongoing development of serial crystallography approaches to enable powerful interrogations of functional motions in PTP1B and other systems, including time-resolved X-ray crystallography with on-chip mixing (Mehrabi et al. 2020) or photoactivation via caged compounds (Monteiro et al. 2021) at synchrotrons or XFELs. Such experiments may enjoy synergy with complementary approaches to interrogate protein motions such as crystal diffuse scattering (Wall, Wolff, and Fraser 2018; Meisburger, Case, and Ando 2020) and solution hydrogen-deuterium exchange mass spectrometry (Kaltashov, Bobst, and Abzalimov 2009).

## Methods

### Protein expression and purification

PTP1B was expressed and purified as reported previously (Keedy et al. 2018). In brief, we used a “wildtype” PTP1B known as WT* containing the C32S/C92V double mutation, residues 1-321, in a pET24b vector resistant to kanamycin. WT* transformed BL21 *E. coli* colonies were selected against LB + kanamycin plates, and used to inoculate 5 mL starter cultures of LB + kanamycin (1 mM final), grown overnight at 37°C with shaking. Starter cultures were subsequently used to inoculate 1 L growth cultures of LB + kanamycin (1 mM final), induced with 100 μM IPTG, and grown for a further 4 hours at 37°C. Cell pellets were harvested via centrifugation and stored at −80°C until needed, or immediately sonicated (on ice) for 10 min, 10 sec on/off at an amplitude of 50%.

PTP1B WT* was purified initially via cation exchange with an SP FF 16/10 HiPrep column (GE Healthcare Life Sciences) in a lysis buffer (100 mM MES pH 6.5, 1 mM EDTA, 1 mM DTT), using a NaCl gradient (0-200mM), with protein eluting around 200mM NaCl. Size exclusion chromatography was subsequently performed using a Superdex S75 size exclusion column (GE Healthcare Life Sciences) in crystallization buffer (10 mM Tris pH 7.5, 0.2 mM EDTA, 25 mM NaCl, 3 mM DTT). Purity was assessed using SDS-PAGE, and pure and contamination free.

### Crystallization

WT* PTP1B was used at 40 mg/mL and drops were set in 96 well plates using an SPT Labtech Mosquito Xtal3, in a ratio of 0.1 μL protein + 0.1 μL well solution (0.1 M MgCl_2_, 0.1 M HEPES pH 7.0, 12-14.5% PEG 4000), incubated at 4°C. Crystals grew within 24 hrs and continued growing for a few more days, reaching final sizes of ~50-100 μm.

### X-ray data collection

Samples were loaded onto the MiTeGen SSX sample supports as reported (Illava et al. 2021), and discussed here in brief. PTP1B crystals and sample supports were placed into a humidified glove box (>97% relative humidity), in order to prevent crystals from drying out on the support during the vacuum loading process. Sample supports were seated within a vacuum port and 3-5 μL of PTP1B crystals in mother liquor were applied to the support surface. A light vacuum was applied to remove excess mother liquor from the support, with the support then sealed using mylar film of thickness 2.5 μm.

Data from PTP1B crystals were collected at room temperature on the ID7B2 (FlexX) beamline at Macromolecular X-ray science at the Cornell High Energy Synchrotron Source (MacCHESS), Ithaca, NY. Crystal-loaded MiTeGen SSX sample supports were mounted on the ID7B2 endstation goniometer, and rastered through the X-ray beam. The sample support was rastered in steps of 20 μm between likely crystal positions. At each such position, 6 data frames were collected, then 4 blank frames were collected during transit to the next position. The 6 data frames consisted of either 0.2° or 0.5° oscillations depending on the chip, for a total oscillation of either 1.2° or 3.0° collected per wedge. Initial data quality and resolution limits were assessed at ID7B2 using the ADX software suite. The photon flux was 10^11^ ph/s, allowing the calculation of an estimated diffraction weighted dose (DWD) per crystal of < 35.1 kGy using RADDOSE-3D (Bury et al. 2018), which is less than the 0.38 MGy = 380 kGy limit proposed for RT SSX (la Mora et al. 2020). We ensured to keep the crystals centered during data collection, and due to the inherent nature of crystal-to-crystal variation in serial crystallography, we report the estimated DWD as an average value. All data collection statistics are reported in **Table 1**.

### Crystallographic data processing

Diffraction data were processed using both XDS (Kabsch 2010) and DIALS (Winter et al. 2022).

For the XDS pipeline, inputs were created using a custom script that ran the generate_XDS.INP script for each wedge in each chip. As noted above, each wedge consists of 10 frames, with the first 6 used for processing, and the last 4 blank frames excluded by the script. The script generated XDS.INP files and ran the parallelized version of XDS (xds_par) for each wedge. See **Fig. 1** for the list of steps run by xds_par. Space group P 31 2 1 was explicitly enforced. Subsequent scaling and merging of integrated intensities across all wedges across all used chips was performed using XSCALE. XDSCONV was used to convert the unmerged XSCALE.HKL file to a merged.hkl file, and then to a merged .mtz file, which was used for subsequent processes.

For the DIALS pipeline, data was imported using dials.import followed by spotfinding using dials.find_spots. A custom Python script was used to identify wedges as sets of consecutive frames with at least 20 identified diffraction spots, excluding blank images (see above), with the result stored in a single .expt and .refl file. Data for all wedges was indexed using dials.index with the flags joint_index=false and beam.fix=all detector.fix=all. Results were refined using dials.refine, and dials.split_experiments was run using refined.* files to create split_*.refl and split_*.expt files for individual wedges. Within a separate directory for each wedge, dials.integrate was run. Successfully integrated wedges were scaled and merged using xia2.multiplex (Gildea et al. 2022), using flags to impose the space group (symmetry.space_group=P3121), completeness (min_completeness=0.95) and resolution (d_min=2.40). The resulting merged .mtz file was used for subsequent processes.

For **Table 1** and **Table 2**, Wilson B values were obtained by running phenix.table_one with the merged data, and some statistics not initially provided by XDS were obtained by running phenix.merging_statistics with the final unmerged data from XSCALE.

### Structure refinement and modeling

The structure was solved using molecular replacement via Phaser (McCoy et al. 2007) using PDB ID 1t49 (with waters and the allosteric inhibitor excluded) as the search model. Iterative rounds of refinement were performed using phenix.refine (Adams et al. 2010) and Coot (Emsley et al. 2010), with model quality validated using MolProbity (Chen et al. 2010; Williams et al. 2018). Data reduction and refinement statistics are reported in **Table 2**. Figures were prepared using PyMOL version 2.5 (Schrödinger, n.d.) via .pml scripting.

## Data availability

Model coordinates and structure factors at the Protein Data Bank with PDB ID 8du7. Raw diffraction images are available on request and will soon be made available publicly.

## Acknowledgments

DAK is supported by NIH R35 GM133769.

This work is based upon research conducted at the Center for High Energy X-ray Sciences (CHEXS), which is supported by the National Science Foundation under award DMR-1829070, and the Macromolecular Diffraction at CHESS (MacCHESS) facility, which is supported by award 1-P30-GM124166-01A1 from the National Institute of General Medical Sciences, National Institutes of Health, and by New York State’s Empire State Development Corporation (NYSTAR).

We thank Irina Kriksunov and Marian Szebenyi for assistance with data collection and initial data processing trials.

We thank Kay Diederichs, Graeme Winter, Dominika Borek, and the rest of the participants from the 2022 CCP4/APS School in Macromolecular Crystallography for help with data processing with XDS and DIALS.

