## Supplemental Tables & Figures for "Room-temperature serial synchrotron crystallography of apo PTP1B"

| | Chip 1 | Chip 2 | Chip 3 | Chip 4 | Chip 5 | Chip 6 | Total |
|---|---|---|---|---|---|---|---|
| XDS | 22 | 18 | 18 | 14 | 20 | 37 | 129 |
| DIALS | 16 | 19 | 16 | 7 | 23 | 37 | 118 |

**Supplemental Table 1: Successfully integrated wedges per chip.**

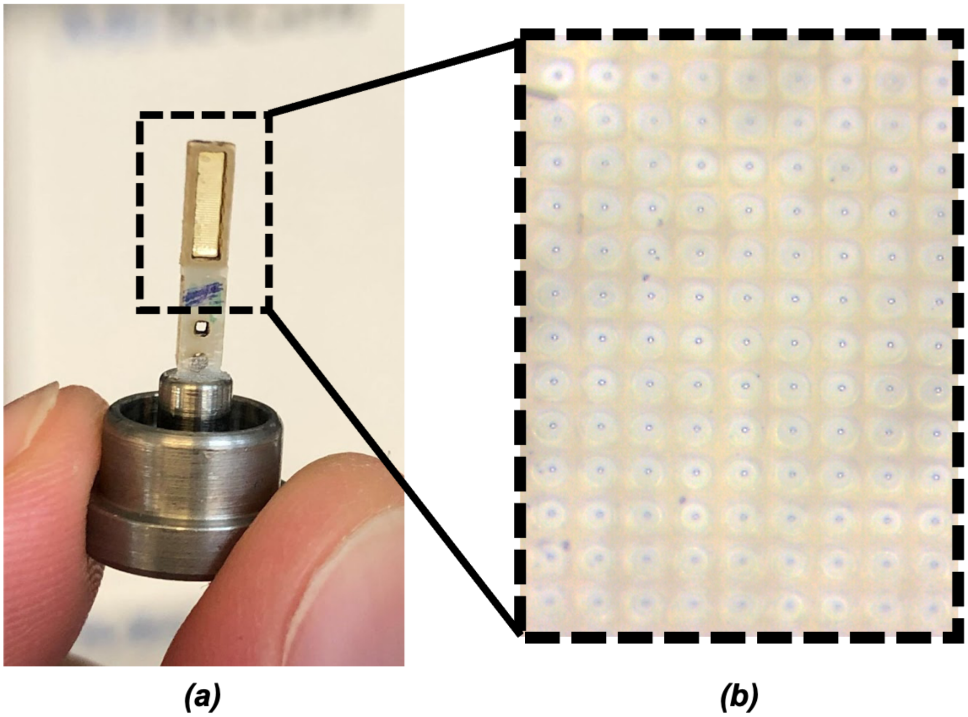

**Supplemental Figure 1: Serial sample support chip used for RT SSX of PTP1B.**
*(a)* Photograph of an example chip with its goniometer-compatible base.
*(b)* Micrograph of crystal-loading facets on the face of a chip.

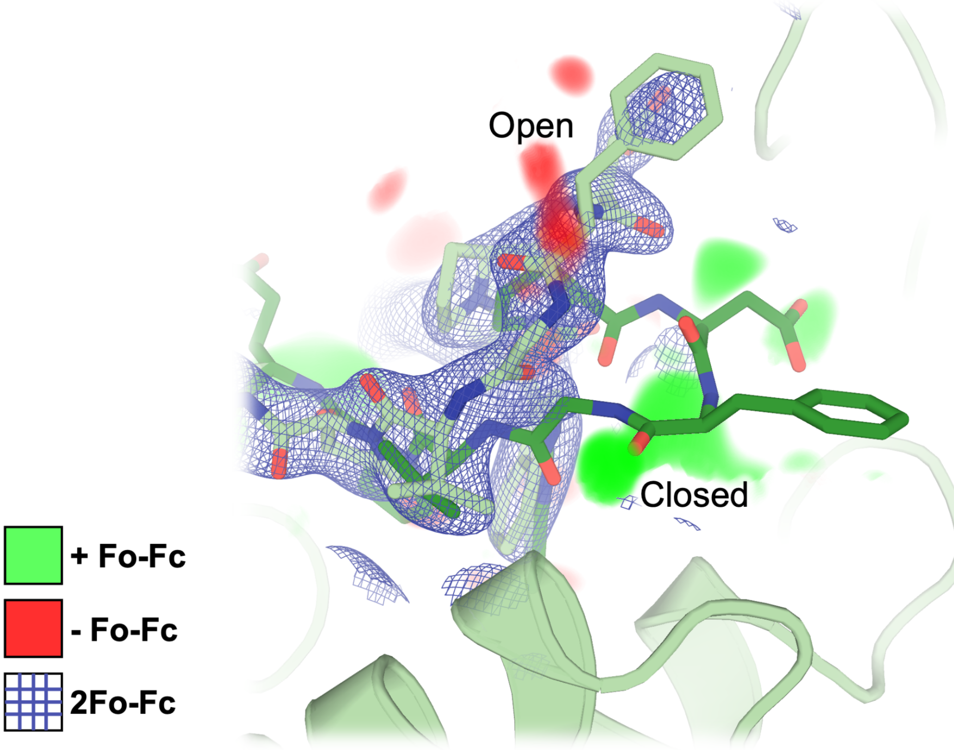


**Supplemental Fig. 2: A dual-conformation model of the WPD loop lacks electron density support.**
2Fo-Fc density at 1 σ and Fo-Fc density at +/- 3.0 σ.

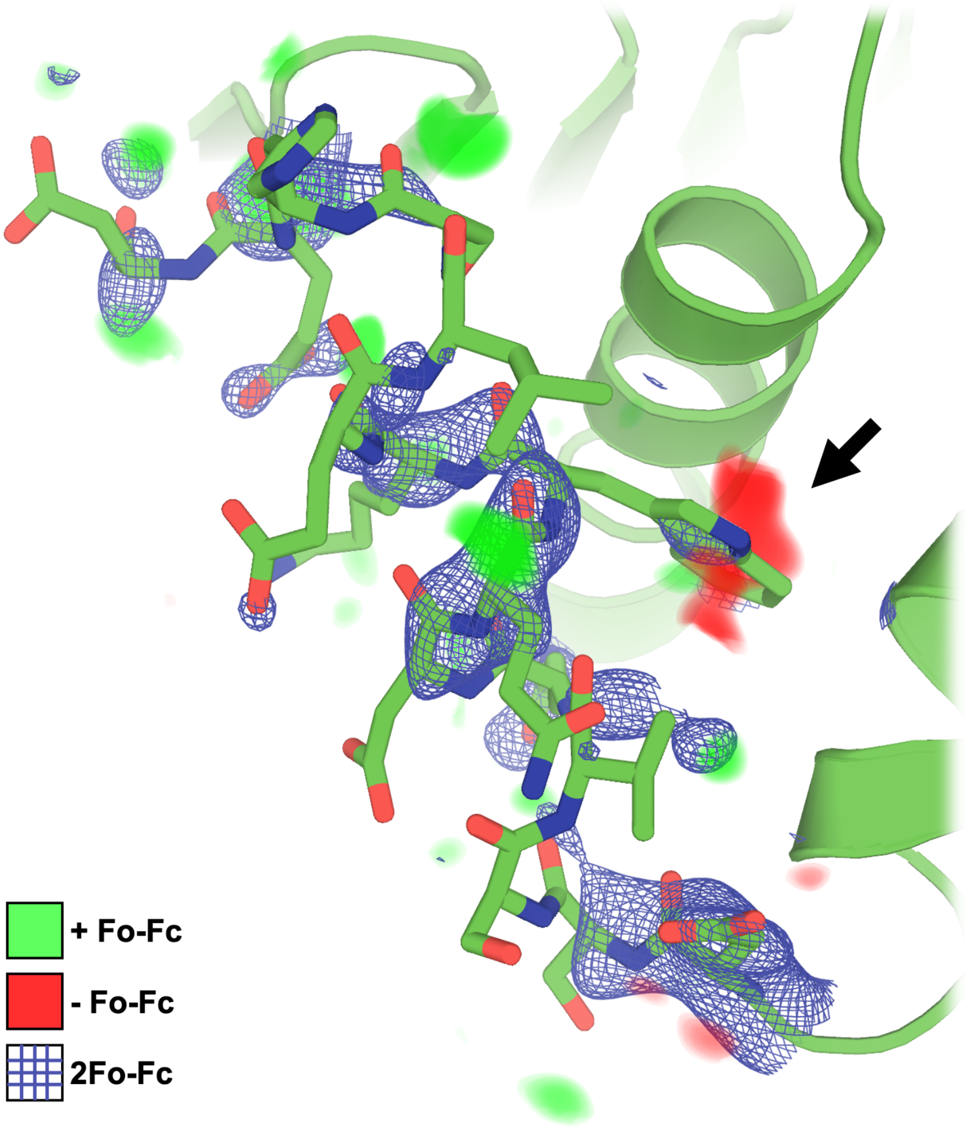


**Supplemental Fig. 3: A model with the ordered α7 helix modeled at partial occupancy has only minimal electron density support.**
2Fo-Fc density at 1 σ and Fo-Fc density at +/- 3.0 σ. Note the negative Fo-Fc density peak for the Trp291 "anchor" when it is modeled as occupying the allosteric BB site (arrow).

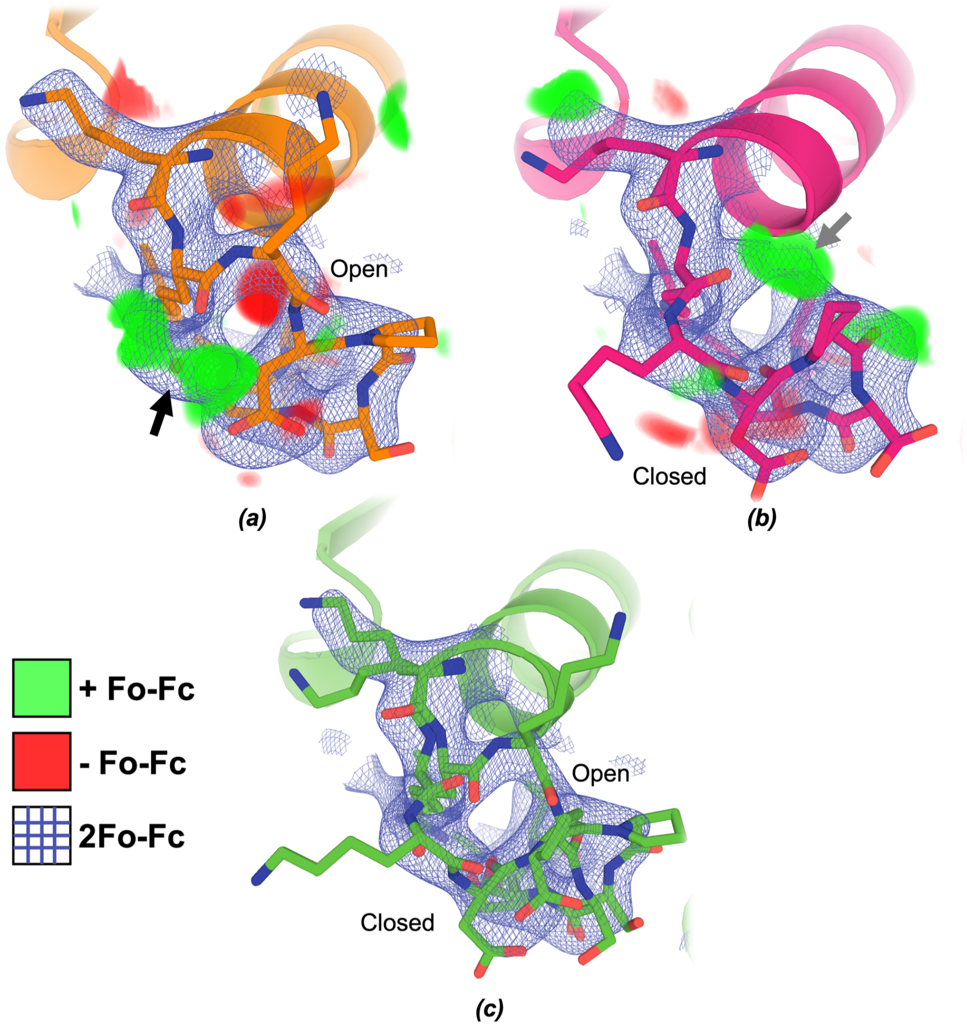


**Supplemental Fig. 4: Omit maps confirm that Loop 16 has alternate conformations.**
2Fo-Fc (1 σ) and Fo-Fc (+/- 3 σ) maps are shown for models with individual states of Loop 16 in the L16 site omitted.
*(a)* Closed state omitted. Positive Fo-Fc density (green) is apparent despite the presence of a modeled water in the 2Fo-Fc map (black arrow), indicating further model building is required.
*(b)* Open state omitted. Positive Fo-Fc density is present for the open L16 conformation (gray arrow), further indicating the model should be built sampling both the open and closed L16 states.
*(c)* Final model including both states as alternate conformations (same as **Fig. 2d**).
